# Virucidal activity of Nasaleze^®^ Cold & Flu Blocker and Nasaleze^®^ Travel in cell cultures infected with human pathogenic coronavirus 229-E

**DOI:** 10.1101/2021.09.23.461483

**Authors:** N Hunt, L Suleman, PD Josling, TA Popov

## Abstract

This *in vitro* study determined the anti-viral efficacy of a unique blend of powder cellulose supplemented with powdered garlic extract (PGE) and a signalling agent. The composition, presented as Nasaleze^®^ Cold & Flu Blocker/Nasaleze^®^ Travel, was assessed against Human Coronavirus 229E, CoV 229E {ATCC VR-740} in an *in vitro* experiment. The test substance was used at sub-optimal dosing levels to explore its prevention and treatment capabilities. The virucidal activity of this novel formulation was measured at 48, 72 and 112 hour periods after incubation. Results showed strong reductions in viral titre of Coronavirus 229E compared to a control, while no toxicity to human cells from the test formulation was noted. The extract Nasaleze^®^ Cold/Travel showed potential to be used as a therapeutic and preventive agent.

The data reconfirms the established anti-viral activity of this formulation acting as a barrier preventing the virus from accessing the nasal mucosa and disrupting its replication.^1,2,3^

## Introduction

The **COVID-19 epidemic in the United Kingdom** is part of the worldwide pandemic of coronavirus disease 2019 (COVID-19) caused by severe acute respiratory syndrome coronavirus 2 (SARS-CoV-2). The virus reached the country in late January 2020. As of 30^th^ August 2020, there have been 334,467 confirmed cases and 41,499 deaths of confirmed cases, the world’s fourth-highest death rate. Worldwide more than 27 million cases and over 891,000 deaths have been recorded, with the United States, Brazil and India recording the highest number of cases. More than 90% of those dying had underlying illnesses or were over 60 years old.

In March 2020, the UK government imposed an order, dubbed “Stay Home, Protect the NHS, Save Lives”, banning all non-essential travel and contact with people outside one’s home (including family and partners), and shutting almost all schools, businesses, venues, facilities, amenities and places of worship. Those with symptoms, and their households, were told to self-isolate 14 days, while those at higher risk due advanced age and accompanying comorbidities were told to shield themselves. People were told to keep apart in public. Police were empowered to enforce the measures, and the Coronavirus Act 2020 gave the government emergency powers not used since the Second World War.

The lengthy restrictions severely damaged the UK economy, lead to millions of job losses, worsened mental health and suicide rates, and caused “collateral” deaths due to isolation and decline of living standards.

In recent years, a number of countries in East and South-East Asia including China had seen an outbreak of various types of infectious flu including SARS Cov 1, MERS, H5N1 avian flu and now Coronavirus described as COVID-19. The infection mainly affected poultry (chickens and ducks) or bats, which were then wiped out in their hundreds of thousands.

The highly pathogenic avian flu virus arrived in Russia in July 2005 and to date the H5N1 flu virus has been recorded in many parts of the Russian Federation: in Western Siberia, in the Urals and in the Astrakhan province. This prompted the conduct of *in vitro* tests using Nasaleze^®^ Cold/Travel which proved very successful at both destroying H5N1 and preventing its replication in human cell lines. This data was published in the European Journal for Nutraceutical Research^3^. Subsequently, a similar *in vitro* test against Coronavirus 229E which is part of the corona virus and common cold virus families with similar characteristics and structures were carried out. With the agent picked for this evaluation we already had a history of success in controlling the viral agent H5N1, so our aim was to check if this Nasaleze^®^ Cold/Travel formulation could be successful in both reducing viral load of Covid 229E and preventing its replication.

## Material and methods

The test viral organism chosen was Human Coronavirus 229E and the utilised cell type was MRC-5. This Medical Research Council cell strain 5 is a diploid human cell culture line composed of fibroblasts, originally developed from lung tissue.

### Cell maintenance and assay set-up

MRC-5 cells were used as the host cell line for human coronavirus 229E (CoV 229E) propagation. MRC-5 cells were maintained in Eagle’s Minimum Essential Medium (EMEM) supplemented with 20% Foetal Bovine Serum (FBS) and 1% penicillin-streptomycin (complete EMEM) at 37 ± 2 °C and 5% CO2. In preparation for the cytotoxicity screening and anti-viral assays, MRC-5 cells were seeded into 24 well plates at 1.0 × 105 cells/mL and incubated at 37 ± 2 ⍰C and 5% CO2 for 24 hours, or until they reached 80-90% confluency. In preparation for tissue culture infectivity dose 50 (TCID50) testing, MRC-5 cells were seeded into 96 well plates at 2 × 105 cellsmL-1 and incubated at 37 ± 2 °C and 5% CO2 for 24 hours.

### Phase 1: Checking for potential cytotoxic effects of the Nasaleze^®^ Cold/Travel formulation on the selected MRC-5 cell line

Nasaleze^®^ Cold/Travel was diluted to 3.2 mg/0.1mL, 6.4 mg/0.1mL and 12.8 mg/0.1mL in EMEM supplemented with 2% FBS and 1% penicillin-streptomycin (assay medium). Complete EMEM was aspirated from the test plates and 100 μL of each test concentration was added to duplicate wells. Following a 10-minute incubation period at 20 ± 2 °C an additional 400 μL of assay medium was added to the test wells. Plates were incubated for 24 hours at 37 ± 2 ⍰C and 5% CO2. Following incubation, visual scoring was performed on a scale of 0 to 4 according to ISO 10993-5 guidelines (Table 1). Cytotoxic effects were assessed based on a variety of morphological changes to the MRC-5 cells such as cell rounding, detachment and cell lysis.

**Table 1.** Cytotoxicity visual scoring and reactivity classifications.

| Visual Score | Cells with cytotoxic effects (%) | Reactivity classification |
| --- | --- | --- |
| 0 | 0 | None |
| 1 | 0 – 20 | Slight |
| 2 | 20 – 50 | Mild |
| 3 | 50 – 70 | Moderate |
| 4 | 70 – 100 | Severe |

### Phase 2: Assessment of the preventative and virucidal capabilities of Nasaleze^®^Cold/Travel

MRC-5 cells were treated with Nasaleze^®^ Cold/Travel according to two methods to determine the preventative and treatment capabilities of the formulation. The assays were performed in 24-well plates utilising duplicate wells for each experimental condition.

### Preventive treatment of MRC-5 cells using Nasaleze^®^ Cold/Travel before infection with high and low doses of human coronavirus 229E

To assess the preventative capabilities of Nasaleze^®^ Cold/Travel against CoV 229E, MRC-5 cells were pre-treated with 3.2 mg of the formulation for 10 minutes before infection with CoV 229E multiplicity of infections (MOIs) of 1 (high dose) and 0.01 (low dose). Complete EMEM was aspirated from the test plates and washed once in Dulbecco’s phosphate buffered saline (DPBS) before application of 3.2 mg Nasaleze^®^ Cold/Travel in 100 μL assay media. Following a 10 minute incubation at 20 ± 2 °C, cells were inoculated with 100 μL CoV 229E, pre-diluted to achieve the high and low MOI infection, and incubated at 35 ± 2°C and 5% CO2 for 30 minutes. Infected cells were then supplemented with an additional 300 μL of assay medium and incubated at 35 ± 2 °C and 5% CO2 for four days. The cytopathic effect (CPE) of the virus on the MRC-5 cells was scored on days 2, 3 and 4 to the criteria described in Table 1. On days 3 and 4, 100 μL of media was harvested from each well to determine the viral titre before replacing with 100 μL of fresh assay medium. Harvested samples were stored at −80 °C until required for viral titre determination. It should be noted that 3.2mg of the test substance is sub optimal dosing and represents only 1 puff into only 1 nostril, whereas the product instructions indicate multiple dosing into BOTH nostrils to prevent or treat any type of airborne infection.

### Treatment of human coronavirus 229E infected MRC-5 cells with Nasaleze^®^ Cold/Travel

To assess the treatment capabilities of Nasaleze^®^ Cold/Travel against CoV 229E, MRC-5 cells were first infected with high and low CoV 229E MOIs, 1 and 0.01 respectively, before treatment with the formulation. Complete EMEM was aspirated from the test plates and washed once in DPBS before being inoculated with 100 μL of pre-diluted CoV 229E to achieve high and low MOI infections and incubated at 35 ± 2 °C and 5 % CO2 for 30 minutes. Following incubation, viral inoculum was removed and a sub optimal 3.2 mg dose of Nasaleze^®^ Cold/Travel in 100 μL assay media was added to the cells and incubated for 10 minutes at 20 ± 2 °C to allow the formation of the gel barrier. Treated cells were then supplemented with an additional 300 μL of assay medium and incubated at 35 ± 2 °C and 5% CO2 for four days. The CPE of the virus on the MRC-5 cells was scored on days 2, 3 and 4 to the criteria described in Table 1. On days 3 and 4, 100 μL of media was harvested from each well to determine the viral titre before replacing with another 100 μL of fresh assay medium. Harvested samples were stored at −80 °C until required for viral titre determination.

### Viral infectivity quantification by TCID50

To determine the viral titre of harvested samples, 10-fold serial dilutions were performed in assay medium. Medium was aspirated from the wells of the cell plate and cells were washed with DPBS. One hundred microlitres of each dilution of the samples were added to the corresponding test wells. Test plates were incubated at 35 ± 2 °C and 5% CO_2_ for 7 days. There were four replicate wells for each test condition. After incubation, viral CPE was determined using an Olympus CK2 inverted microscope. The viral titre was calculated using the Spearman-Kärber method.

## Results

### Phase 1: MRC-5 cytotoxicity screen

There was no observable cytotoxicity in MRC-5 cells exposed to Nasaleze^®^ Cold/Travel following a 24-hour contact time (Table 2). When visual scoring was performed, the gel barrier formed by Nasaleze^®^ Cold/Travel was visible on top of the cell monolayer. Additionally, a residue was visible on treated cells (Appendix I).

**Table 2.** Cytotoxicity of Nasaleze^®^ Cold/Travel using visual scoring.

| Treatment | Visual score | Reactivity classification |
| --- | --- | --- |
| Nasaleze<sup>®</sup> Cold/Travel | 0 | No cytotoxicity |

### Preventive treatment of MRC-5 cells using Nasaleze^®^ Cold/Travel before infection with coronavirus 229E

#### Cytopathic effect of CoV 229E on MRC-5 cells pre-treated with Nasaleze^®^ Cold/Travel

Following a 2, 3 and 4 day or 48, 72 and 112 hours incubation period, the CPE of the test plate was scored (Chart 1). Representative images of the CPE observed are presented in Appendix II. Duplicate cells treated with Nasaleze^®^ Cold/Travel with a high MOI of CoV 229E showed slight CPE on day 2 and severe CPE on days 3 and 4. Duplicate cells treated with Nasaleze^®^ Cold/Travel with a low MOI of CoV 229E showed no CPE on day 2 and moderate CPE on days 3 and 4

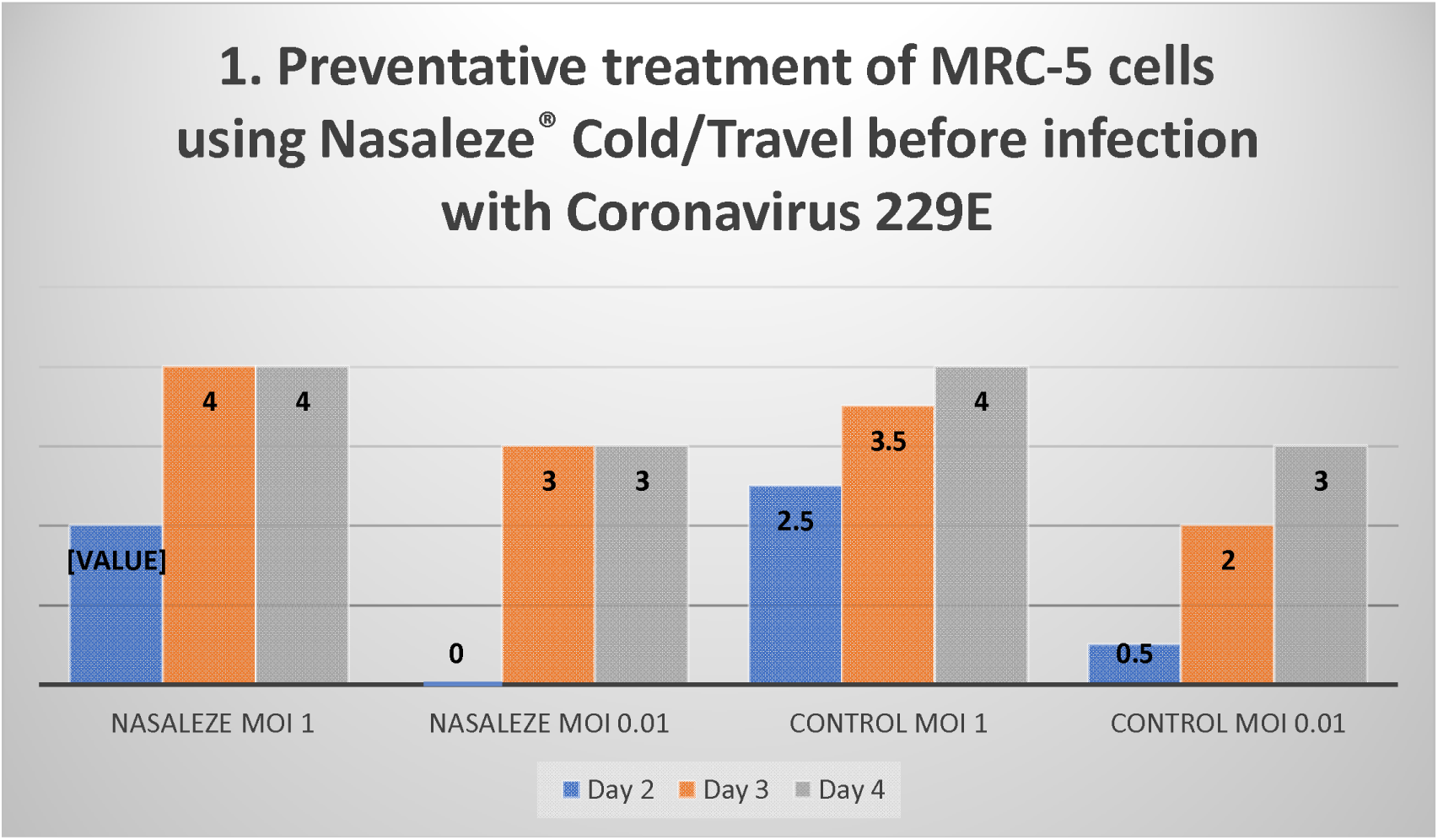

Following a 3 and 4 day incubation period with a high MOI of CoV 229E the negative control resulted in an average viral titre of 5.82 ± 0.35 Log_10_TCID_50_/mL and 5.32 ± 0.35 Log_10_TCID_50_/mL, respectively. Pre-treatment of MRC-5 cells with Nasaleze^®^ Cold/Travel resulted in a 2.68 Log_10_TCID_50_/mL and 2.55 Log_10_TCID_50_/mL reduction in viral titre on day 3 and day 4 post-infection, respectively, when compared to the negative control showing an average of 3.14 ± 0.18 and 2.77 ± 0.53 Table 3 and 4 - Chart 2.

**Table 3.** Log TCID50 and Log reduction values for human coronavirus 229E (CoV 229E) following treatment with Nasaleze^®^ Cold/Travel before infection at a high multiplicity of infection and incubated for 3 and 4 days. N/A = not applicable, SD = standard deviation.

| Product | Average Viable CoV 229E ± SD (Log<sub>10</sub>TCID<sub>50</sub>/mL) |  | Log Reduction (Log<sub>10</sub>TCID<sub>50</sub>/mL) |  |
| --- | --- | --- | --- | --- |
|  | Day 3 | Day 4 | Day 3 | Day 4 |
| Negative Control | 5.82 ± 0.35 | 5.32 ± 0.35 | N/A | N/A |
| Nasaleze<sup>®</sup> Cold/Travel | 3.14 ± 0.18 | 2.77 ± 0.53 | 2.68 | 2.55 |

**Table 4.** Log TCID50 and Log reduction values for human coronavirus 229E (CoV 229E) following treatment with Nasaleze^®^ Cold/Travel before infection at a low multiplicity of infection and incubated for 3 and 4 days. N/A = not applicable, SD = standard deviation.

| Product | Average Viable CoV 229E ± SD ($\text{Log}_{10}\text{TCID}_{50}/\text{mL}$) | | Log Reduction ($\text{Log}_{10}\text{TCID}_{50}/\text{mL}$) | |
| --- | --- | --- | --- | --- |
|  | Day 3 | Day 4 | Day 3 | Day 4 |
| Negative Control | $6.02 \pm 0.53$ | $5.39 \pm 0.18$ | N/A | N/A |
| Nasaleze® Cold/Travel | $4.32 \pm 0.35$ | $4.39 \pm 0.18$ | 1.70 | 1.00 |

### Large viral titre

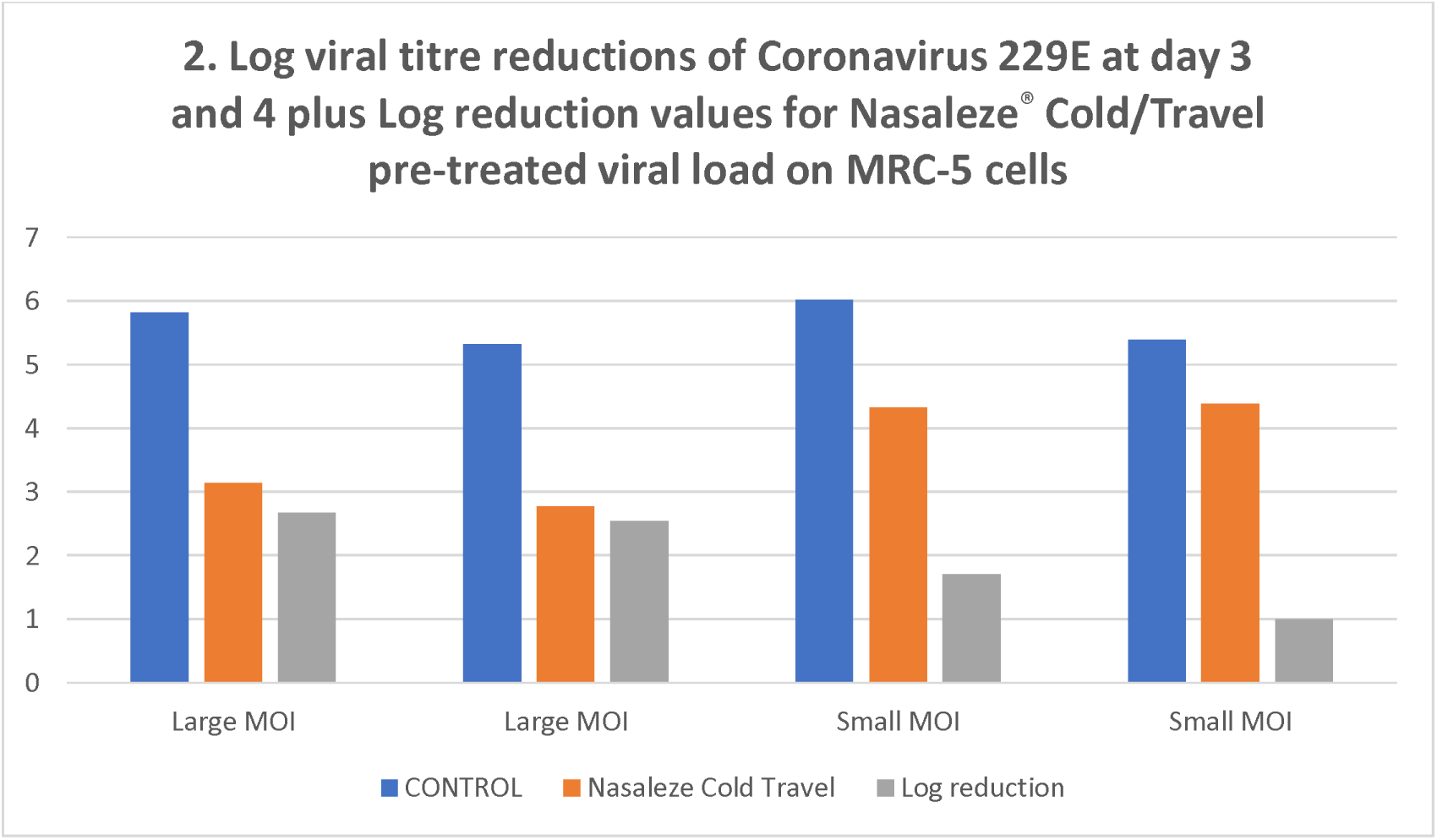

### Small viral titre

Following a 3 and 4 day incubation period with a low MOI of CoV 229E the negative control resulted in an average viral titre of 6.02 ± 0.53 Log_10_TCID_50_/mL and 5.39 ± 0.18 Log_10_TCID_50_/mL, respectively. Pre-treatment of MRC-5 cells with Nasaleze^®^ Cold/Travel resulted in a 1.70 Log_10_TCID_50_/mL and 1.00 Log_10_TCID_50_/mL reduction in viral titre on day 3 and day 4 post-infection, respectively, when compared to the negative control (Table 4, Chart 2).

### Treatment capabilities of Nasaleze^®^ Cold/Travel

#### Cytopathic effect of CoV 229E on MRC-5 cells treated with Nasaleze^®^ Cold/Travel after viral infection

Following a 2, 3 and 4 day incubation period, the CPE of the test plate was scored. Representative images of the CPE observed are presented in Appendix Duplicate cells treated with Nasaleze^®^ Cold/Travel after infection with a high MOI of CoV 229E showed mild CPE on day 2 and severe CPE on days 3 and 4 post-infection. Duplicate cells treated with Nasaleze Cold/Travel after infection with a low MOI of CoV 229E showed no CPE on day 2 and moderate CPE on days 3 and 4 post-infection.

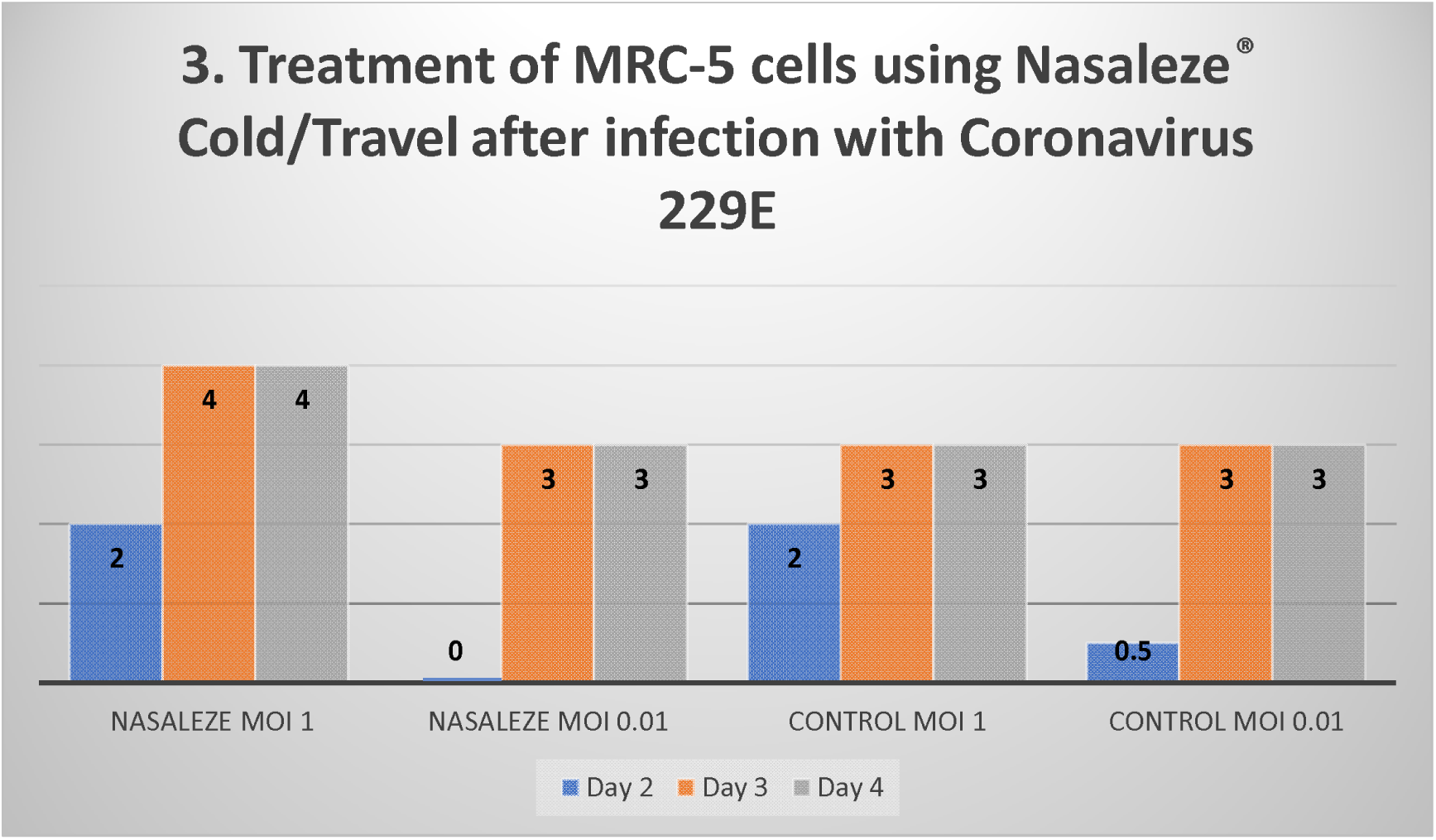

### Viral titration of samples treated with Nasaleze^®^ Cold/Travel Blocker after viral infection Chart 4

Following a 3 and 4 day incubation period with a high MOI of CoV 229E the negative control resulted in an average viral titre of 5.82 ± 0.35 Log_10_TCID_50_/mL and 5.32 ± 0.35 Log_10_TCID_50_/mL, respectively. Treatment with Nasaleze^®^ Cold/Travel after infection resulted in a strong log reduction on days 3 and 4 at 4.75 ± 0.00 and 3.39 ± 0.18 respectively.

Furthermore a 3 and 4 day incubation period with a low MOI of CoV 229E the negative control resulted in an average viral titre of 6.50 ± 0.00 Log_10_TCID_50_/mL and 5.89 ± 0.18 Log_10_TCID_50_/mL, respectively. Treatment with Nasaleze^®^ Cold/Travel after infection resulted in a strong log reduction on days 3 and 4 at 5.75 ± 0.00 and 4.89 ± 0.18 respectively.

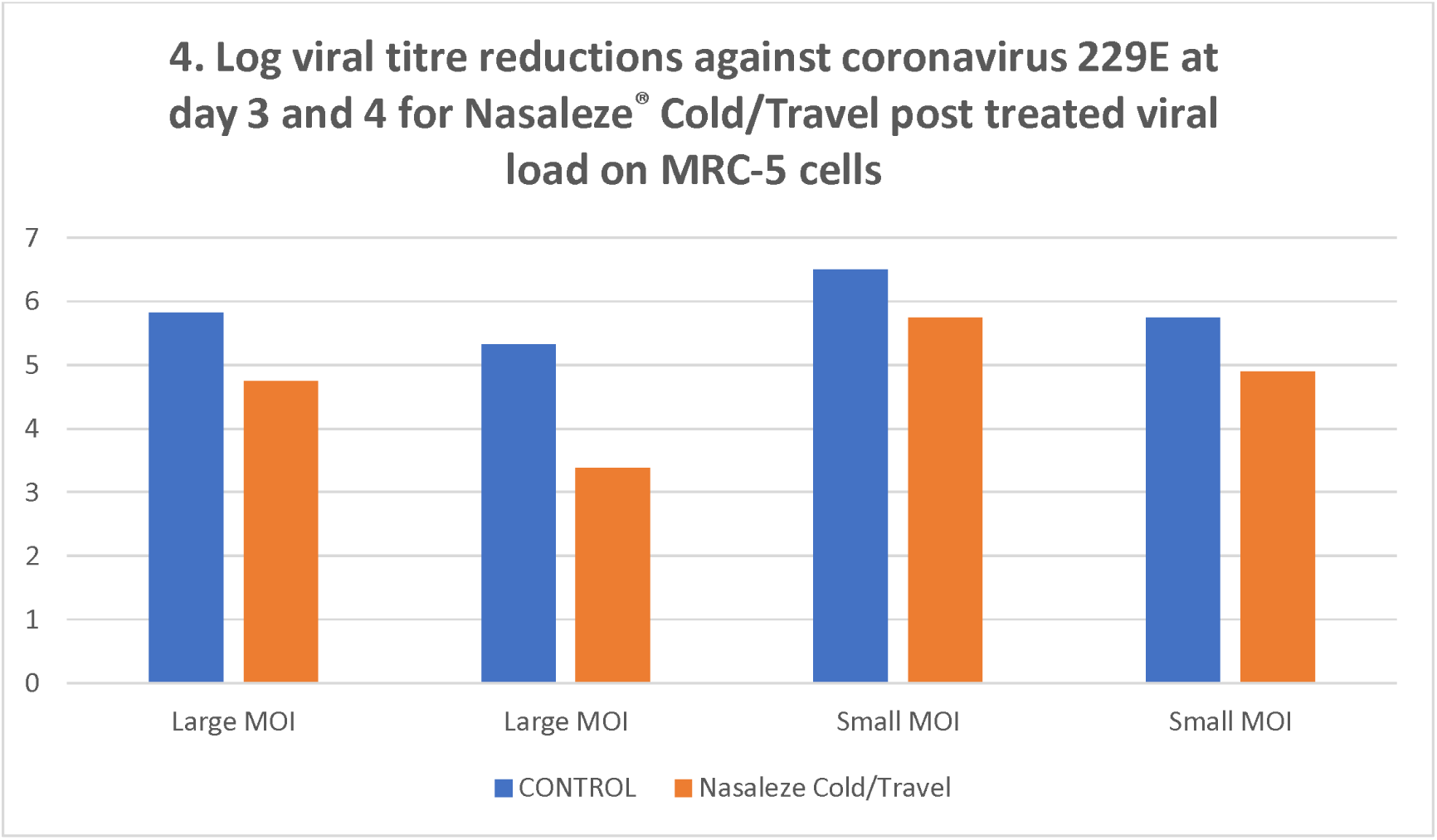

## Discussion

The dissemination of potentially pathogenic viruses increases infection risk in both healthy and immunocompromised individuals. Coronaviruses are enveloped, single stranded RNA viruses responsible for a variety of upper-respiratory tract illnesses in humans. The severity of these illnesses ranges from mild as in common cold to severe acute respiratory syndrome as seen in the recent COVID-19 pandemic. Coronaviruses are thought to be predominantly transmitted through respiratory droplets with some evidence to suggest the virus can remain active on fomites for several days. Interventions, both preventative and curative, are essential to slowing and/or stopping the spread of coronaviruses.

The assessment of interventions against coronavirus surrogate strains allows for the safe evaluation of product efficacy. Coronavirus 299E is structurally and genetically similar to the SARS-CoV-2 virus. Since the COVID-19 pandemic, the Australian regulatory body, Therapeutic Goods Administration (TGA), is the first regulatory body to announce that **Coronavirus 229E as a suitable coronavirus surrogate strain for biocide coronavirus claims**.

Two different experiments were performed to investigate the anti-viral efficacy of Nasaleze^®^ Cold/Travel. In the first experiment, MRC-5 cells were pre-treated with Nasaleze^®^ Cold/Travel before infection with high and low doses of CoV 229E. in the second experiment MRC-5 cells were infected with a high and low dose of CoV 229E before treatment with Nasaleze^®^ Cold/Travel. Treatment with Nasaleze^®^ Cold/Travel did not damage the experimental MRC-5 cell line, but yielded substantial reductions in viral titre indicating a high level of anti-viral potential. Although the reduction in CPE was not large or maintained it should be noted that a sub optimal dose was used representing only one dose into a single nostril, whereas real life clinical data accumulated thus far has shown that a three times daily dose into each nostril can significantly reduce airborne infection.^1,2,3^

Future work could investigate the optimal dosing of Nasaleze^®^ Cold/Travel, simulating the real-life intended use of the product. Additionally, similar experiments could be performed using other respiratory viruses such as influenza, adenovirus and rhinovirus. Finally, as this formulation shows such promise in both preventing and treating viral infection, a 3D primary nasal cell culture model could be considered for use to obtain a more translatable result as well as clinical evaluations in human subjects to add to the existing database for this unique powder cellulose, signalling agent and garlic extract, marketed as Nasaleze^®^ Cold & Flu Blocker and Nasaleze^®^ Travel.

### Key take away points from the report

We asked co-author Peter Josling for his comments on the results…

“This is a very interesting *in vitro* study that clearly shows Nasaleze^®^ Cold & Flu Blocker and Nasaleze Travel are unique active formulations in the fight to both prevent and treat coronavirus infections.

It is clear that pre-treatment reduces viral replication and may therefore stop Coronavirus 229E in its tracks when used at optimum dosing levels.

Even when viral replication is already infecting healthy human cells Nasaleze^®^ Cold/Travel can attack and disable viral replication.

These results are from a SINGLE dose of Nasaleze^®^ Cold/Travel and we would expect multiple doses to be even more effective.

Nasaleze^®^ Cold/Travel do not have any negative cytopathic effect on human cells.

This is a step forward in the prevention and management of coronavirus infection.”

Dr Peter Josling

Herbal Research Centre

## Appendix 1

**Figure A.**
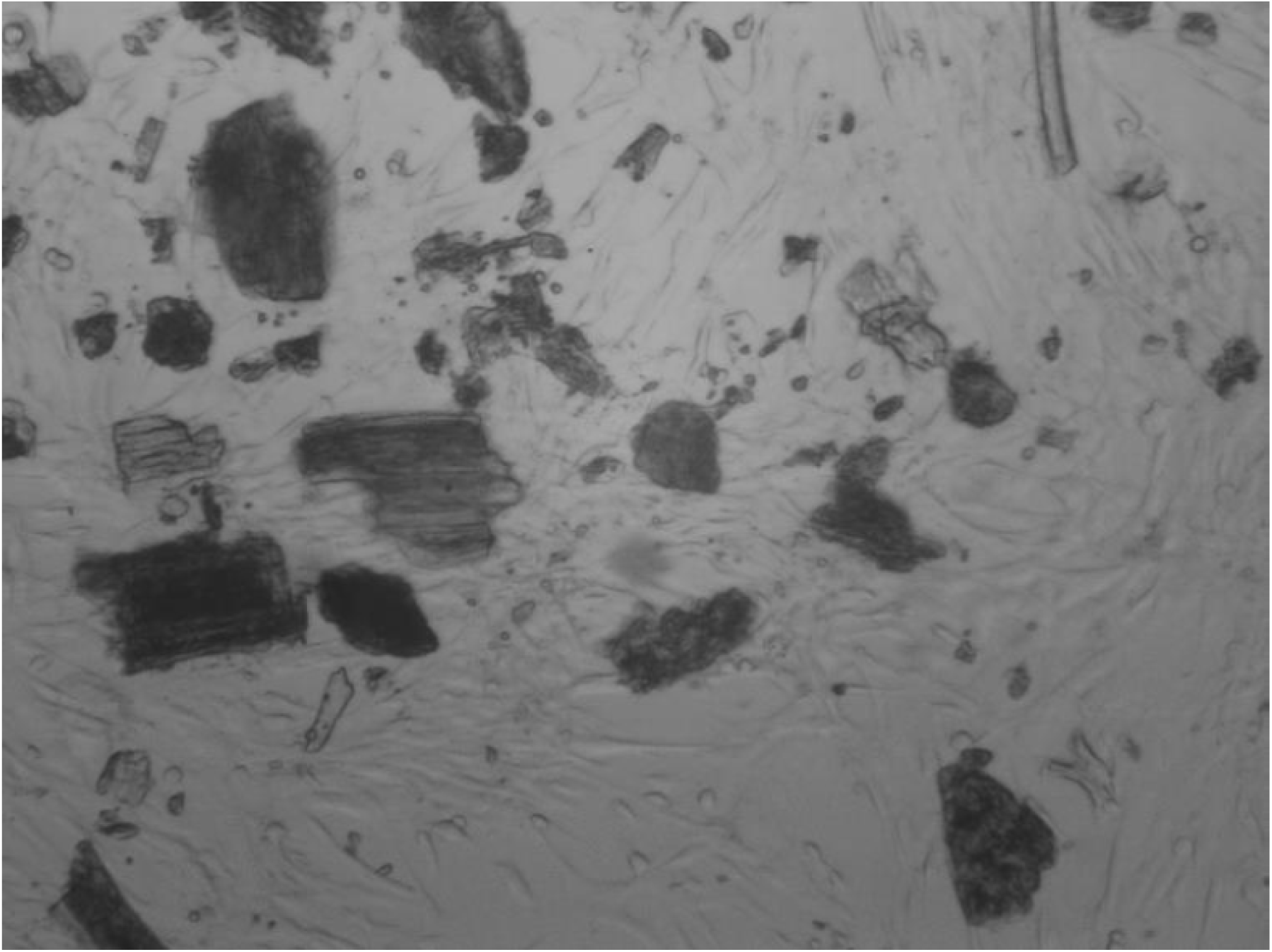
Microscope images showing the residue visible on MRC-5 cells after treatment with Nasaleze^®^ Cold/Travel.

## Appendix II

**Figure B:**
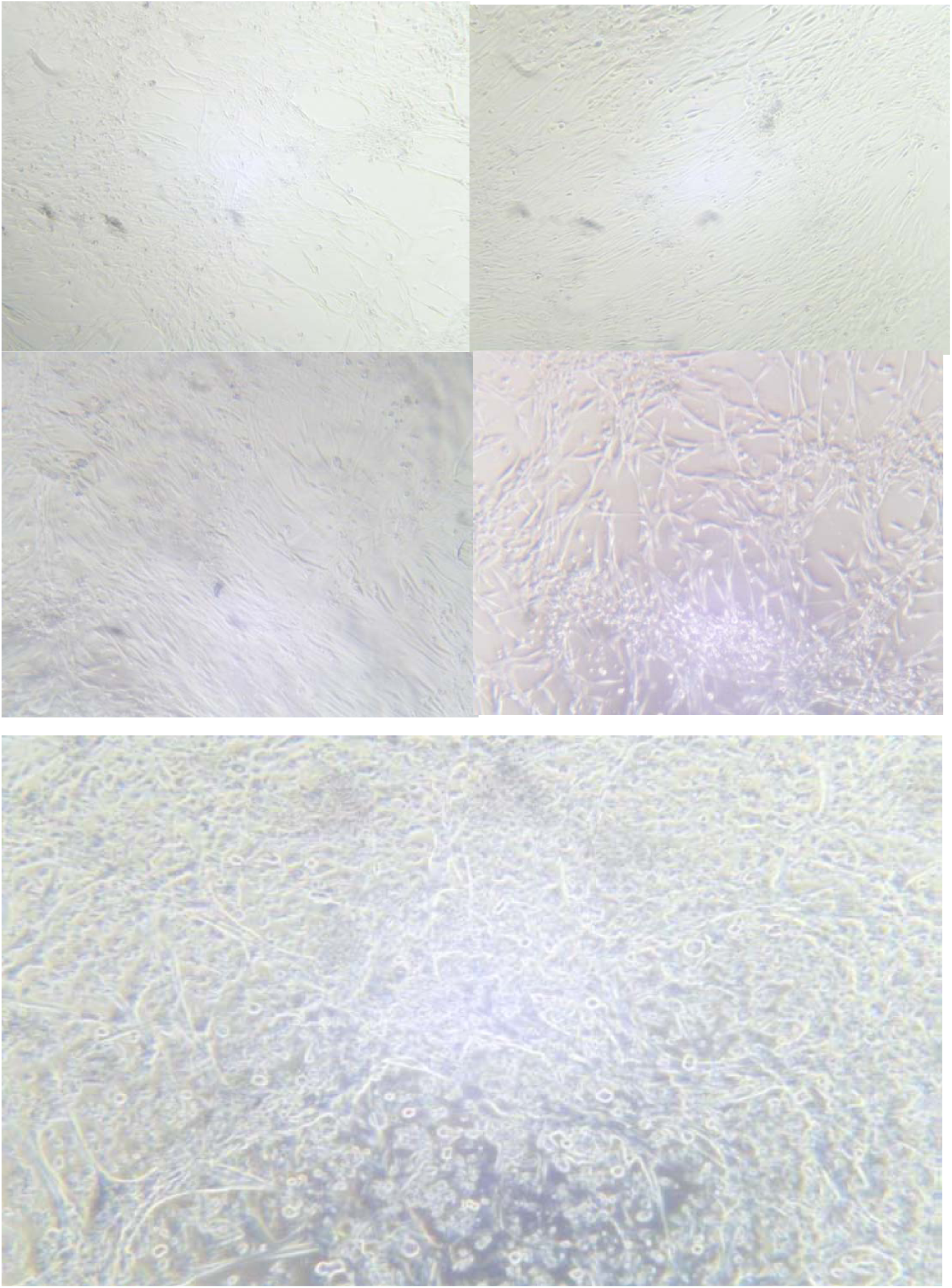
Representative images of cytopathic effect of MRC-5 cells observed throughout the study. Top left = No CPE, Top right = Slight CPE, Middle left = Mild CPE, Middle right = Moderate CPE, Bottom = Severe CPE.

